# Determinants of mutation load in birds

**DOI:** 10.1101/2025.08.18.670172

**Authors:** Fidel Botero-Castro, Jochen B. W. Wolf

**Affiliations:** Department of Evolutionary Biology, Faculty of Biology, LMU Munich, Germany

**Keywords:** Selection, effective population size, recombination, distribution of fitness effects, purging

## Abstract

Many mutations have detrimental effects. The mutation load in a population depends on the efficacy of purifying selection in removing deleterious genetic variation. Here, we estimated the proportion of deleterious mutations segregating in 24 population samples of 19 bird species. Exploiting the conserved avian karyotype with high variation in recombination rate and GC content, we quantified the joint effects of effective population size (*N_e_*), recombination (*r*) and GC-biased gene-conversion (*gBGC*). In agreement with the nearly-neutral theory of molecular evolution, mutation load was substantially higher in populations with small *N_e_*. Purging efficacy increased with recombination rate resulting in more than a two-fold difference of genetic load between large and small chromosomes. GC-biased mutations contributed about one third to the pool of deleterious mutations. Their expected accumulation in regions of high recombination was offset by purging efficacy in large, but not small populations. This study provides insight into how the interaction of evolutionary processes shapes mutation load. It suggests that genetic risk factors in small populations are fueled by *gBGC* and cluster in regions of low recombination.

**Summary:** Harmful genetic mutations can accumulate in populations, affecting individual-and population-level health. The authors analyzed genomic data of 24 populations of 19 bird species to understand the evolutionary determinants of such mutation load. They confirm that small populations tend to experience a heavier mutation burden. Moreover, deleterious mutations accumulate more readily in regions of low recombination and are further elevated by GC-biased gene conversion (*gBGC*), which is offset by purging efficacy in large, but not in small populations. This study suggests that genetic risk factors in small populations are fueled by *gBGC* and cluster in regions of low recombination.

## Introduction

Novel mutations enter a population each generation. Many are neutral, but of those affecting the fitness of its carrier most are deleterious, some are advantageous (Keightley and Lynch 2003). The overall frequency of selective effects is summarized in the genome-wide distribution of fitness effects (DFE, **Figure 1)** reflecting properties of the organism and the distance to its global fitness optimum (Eyre-Walker and Keightley 2007; Lanfear et al. 2014; Chen et al. 2022). Lethal or strongly deleterious mutations are quickly removed from a population by purifying selection, and beneficial mutations are promoted to fixation. Neutral and weakly deleterious mutations, however, can persist for many generations and constitute the bulk of segregating genetic variation. The time they spend in a population depends on the interaction of their fitness effects (*s*) with the effective size of the population (*N_e_*), the population selection coefficient *N_e_s* (Kimura 1983; Ohta 1992; Charlesworth 2009; Akashi et al. 2012). As a consequence, deleterious mutations can rise to high frequencies in populations with small *N_e_* (**Figure 1**).

**Figure 1.**
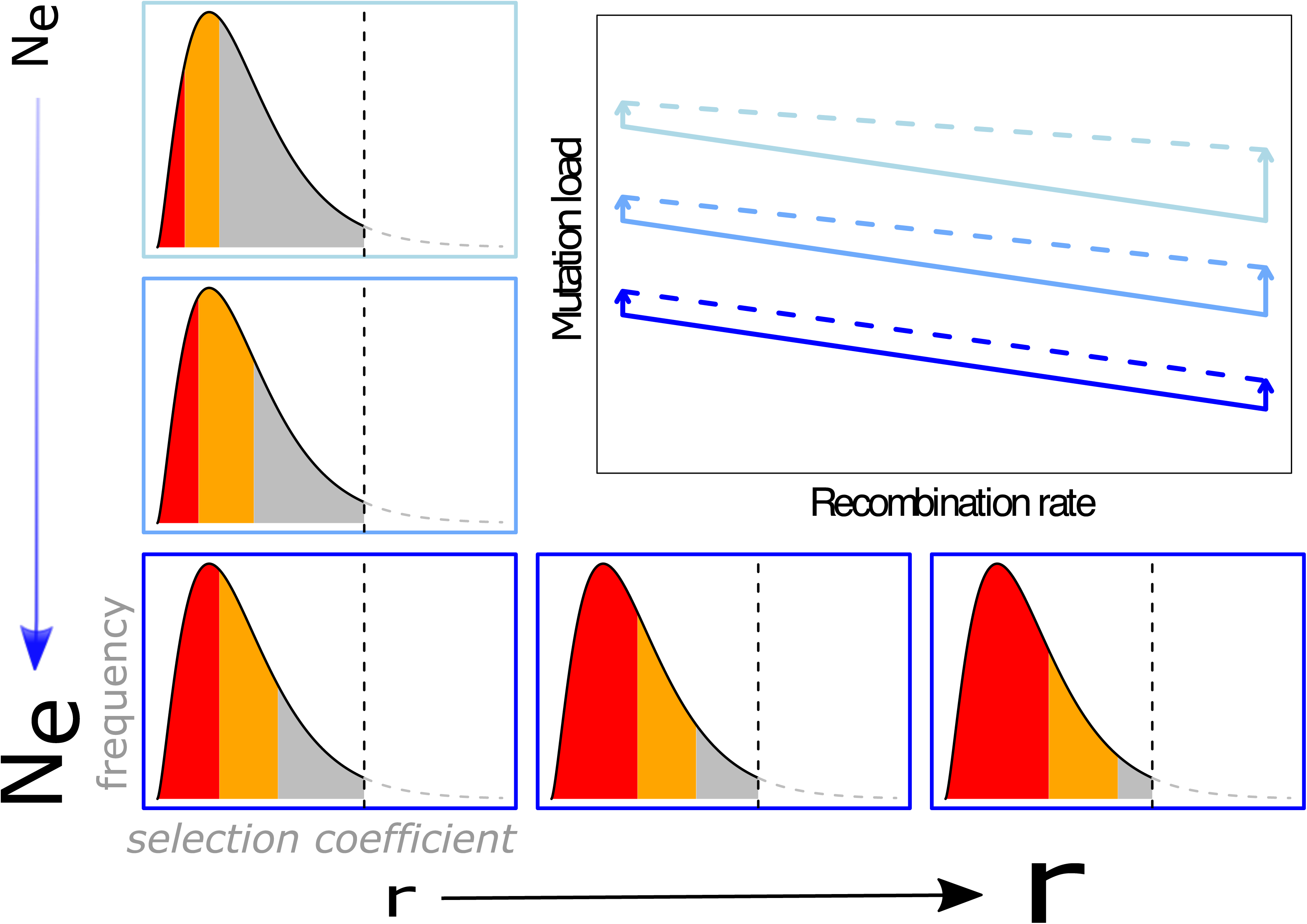
Theoretical considerations on mutation load. The curves outlined in black illustrate a mock example of a genome-wide distribution of fitness effects (DFE) at a given point in the fitness landscape. We here only consider the fraction of mutations with selection coefficient s<0 reducing fitness (left of the vertical dashed line marking *s*=0). The efficacy of selection in removing those crucially depends on the effective population size *N_e_* (left column) and the recombination rate *r* (bottom row) changing the fraction of mutations behaving as strongly deleterious (red: *N_e_s* < -10), deleterious (orange: *N_e_s* ∈ (-10;-1]), or mildly deleterious (grey: *N_e_s* ∈ (-1;0]). The latter are the focus of this study. *Central figure, solid lines:* The y-axis shows the fraction of all mutations falling into the mildly deleterious category *N_e_s* ∈ (-1;0]. This corresponds to the mutation load which is expected to be highest in populations with small N_e_. Mutation load is further expected to be elevated in genomic regions experiencing low recombination rates (x-axis). Line colours correspond to *N_e_* categories small (light blue), intermediate (blue) and large (dark blue). *gBGC* is a recombination-associated process expected to increase mutation load overall, and particularly so in regions of high recombination *(dashed lines*). Difference in slope between solid (without *gBGC*) and dashed lines (with *gBGC*) is expected if the relative rate by which recombination introduces mildly deleterious mutations through *gBGC* differs from the rate by which recombination increases the efficacy of selection removing these. Such difference in relative rates of *gBGC* and purging efficacy may further interact with the effective population size (length of arrows) mediating the overall and/or recombination-dependent contribution of *gBGC* to mutation load.

These considerations of the nearly-neutral theory of molecular evolution are of real-world importance (Dussex et al. 2023). The accumulation of deleterious mutations maintained by mutation-selection balance, the mutation load (Muller 1950), can eventually elevate the risk of population extinction (Lande 1994; Frankham 1998; Soulé and Mills 1998; Agrawal and Whitlock 2012). There is growing empirical evidence that mutation load indeed scales negatively with proxies for *N*_e_ as it has been predicted by theory (Kimura et al. 1963; Lynch et al. 1995). One of such proxies is the size of the geographical range occupied by a given population, which is expected to be directly correlated to the census size of said population, and, ultimately, to *N*_e_ (Peart et al. 2020). Indeed, across a broad taxonomic range, the efficacy of purifying selection tends to be enhanced in species and populations characterized by a large number of effectively breeding individuals (Slotte et al. 2010; Deinum et al. 2015; Chen et al. 2017; Castellano et al. 2018; Galtier and Rousselle 2020). As a consequence, mutation load tends to be increased in small, isolated island populations as compared to populations with larger distribution areas (Kutschera et al. 2020), even when comparing island and continental populations of the same species (Leroy et al. 2021).

While the effect of population size on mutation load has been well studied, other factors have received less attention. Recombination, for instance, is known to modulate the efficacy of selection at the intragenomic scale. In regions of low recombination, Hill-Robertson interference acts as to reduce the local effective population size, thus increasing the probability of deleterious alleles rising to fixation (Hill and Robertson 1966; Felsenstein 1974; Comeron et al. 2008). Similarly, the purging effect of synergistic epistasis between deleterious mutations, though contentious, is influenced by recombination (Agrawal and Whitlock 2012). As a consequence, mutation load is expected to differ within the genome as a function of recombination intensity (Comeron et al. 2008). Empirical evidence comes from exploiting variation in recombination rates within and across chromosomes (Gossmann et al. 2011; Webster and Hurst 2012; Corcoran et al. 2017; Chase et al. 2024). Due to obligate crossover, small chromosomes have high recombination rates per physical unit (cM/Mb) (Hillier et al, 2004; Kawakami et al. 2014, Backström et al 2010, Bascón-Cardozo et al 2023) and are expected to carry the smallest load, whereas large, low-recombining chromosomes are expected to accumulate more deleterious mutations. Non-recombining, uniparentally inherited sex-chromosomes or organellar genomes are predicted hotspots of mutation load and have accordingly received most attention (Charlesworth and Charlesworth 2000; Marais 2007; Warmuth et al. 2022).

Importantly, recombination is also known to be associated with GC-biased gene conversion (*gBGC*), referring to the preferential reparation of double-strand breaks towards G and C alleles, which eventually results in an increase of GC content in highly recombining regions (Galtier et al. 2001; Duret and Galtier 2009). Most of these mutations are prone to deleterious effects but are nevertheless constantly replenished through the segregational bias in regions undergoing high recombination, mimicking the effect of positive selection (Berglund et al. 2009). *gBGC* is thus predicted to counteract efficient purging of weakly-deleterious mutations in regions of high recombination (Bolívar et al. 2018). Yet, it is unclear at which rate these opposing forces operate, in particular, as the strength of *gBGC* also seems to interact with the overall effective population size (Bolívar et al. 2019) (**Figure 1**).

Avian genomes are well suited to study these interacting effects, as they provide the necessary variation in *N_e_* across a stable, yet highly variable, recombination landscape within and between chromosomes (**Figure 2**) (Ellegren 2010). To assess the interaction of 2/4/2026 5:57:00 AMrecombination, effective population size and GC-biased gene conversion, and their influence on mutation load, we collated genome-wide resequencing data across 24 populations from 19 avian species varying in effective population size. We estimated the DFE for individual chromosomes with substantial variation in recombination rate to obtain estimates of the proportion of segregating, mildly-deleterious mutations (*N_e_s* ∈ (-1;0]), serving as proxy for mutation load. According to theory, we expect the following: mutation load should be: 1) higher in populations with low *N_e_*, 2) reduced in regions of high recombination and 3) increased for the mutational spectrum subject to *gBGC* (weak-to-strong mutations: [A,T]->[C,G]). 4) Relative rates of the two recombination-mediated, opposing processes of purging and *gBGC* may differ and be additionally modified by *N_e_* (**Figure 1**).

**Figure 2.**
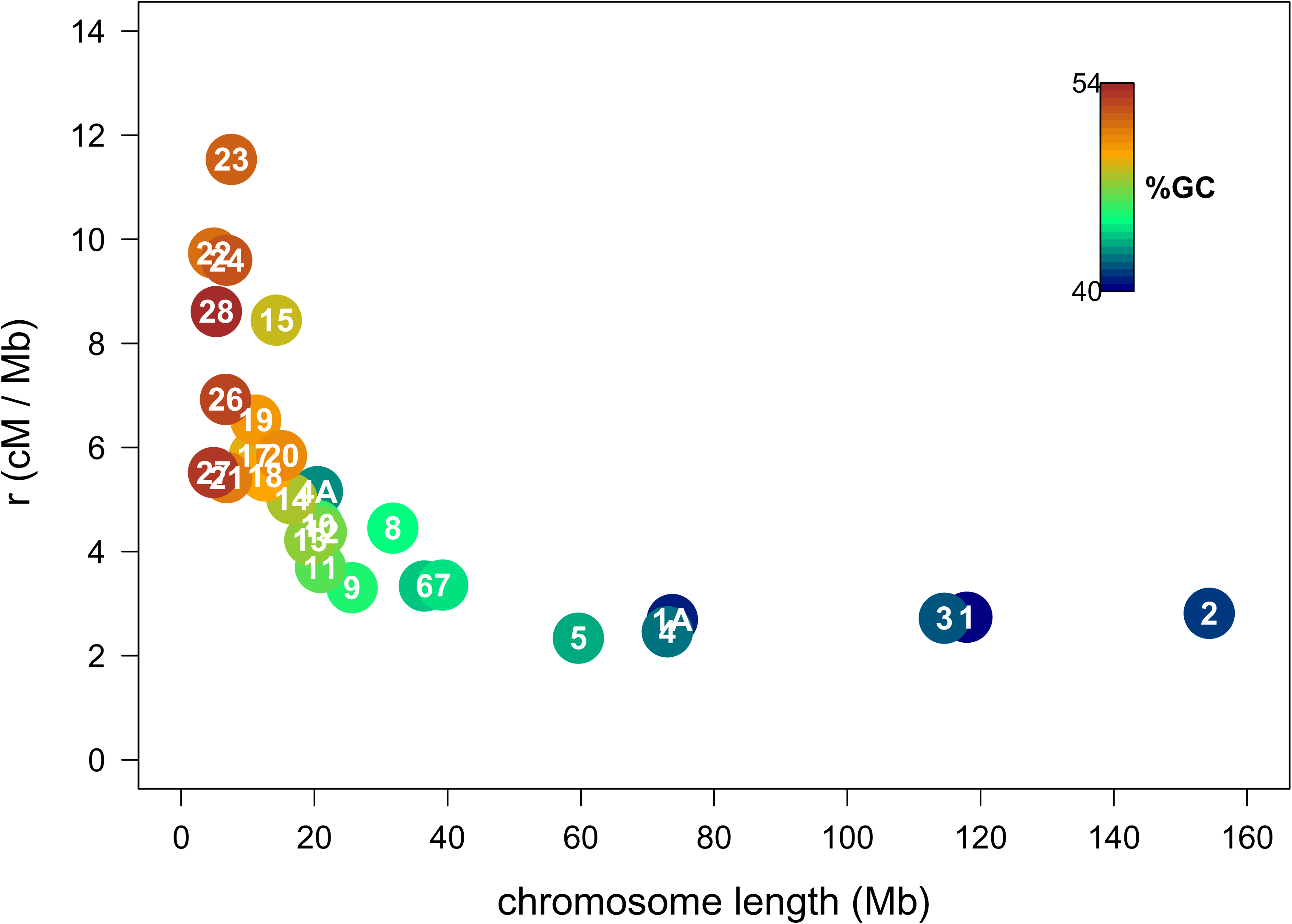
Relationship between recombination rate [cM/Mb], chromosome length [Mb] and GC content [%] in Eurasian crows, *Corvus (corone) ssp*. A near 5-fold difference in recombination rate is closely associated with chromosome length which strongly covaries with GC content (R^2^=0.90, p<0.001). In small, highly recombining chromosomes GC content is increased by up to 15% reflecting an evolutionary history of *gBGC*. The heterogenous recombination landscape is conserved across the avian species investigated in this study.

## Materials and Methods

### Sequence Data

We collated publicly available resequencing data for 443 individuals from 47 populations of 30 passerine (sub)species and utilized *.vcf* files generated by comparable pipelines described in (Leroy et al. 2021) and (Kutschera et al. 2020) (see **Supplementary Table 1** for accession numbers of NCBI’s Sequencing Read Archive). In summary, reads were first mapped to the closest available reference (see **Supplementary Table 1**). After the usual steps of adding read group information, merging bam files per individual, marking duplicates and realignment of indels, both studies used GATK (v3.4.0 in Kutschera et al (2020), v3.7.0 in Leroy et al. (2021)) to call variants using joint genotyping with *GenotypeGVCF*. This resulted in *vcf* files including both variant and invariant sites.

While Leroy et al. (2021) directly proceeded to filter the output of *GATK* on the basis of quality by depth (QD < 2.0), Fisher Strand bias (FS > 60), mapping quality (MQ < 40), MQranksum < 2 or ReadPosRankSum < 2 or Raw Mapping Quality (Raw_MQ < 45,000), Kutschera et al (2020), performed an additional step of base-quality score recalibration (BQSR) before jointly genotyping, using a set of “true” variants resulting from the highest quality variants identified in the intersection of the outputs of *samtools* v1.3 (Li et al. 2009), *FreeBayes* v1.0.2 (Garrison and Marth 2012), and *GATK*’s HaplotypeCaller. Then, after joint genotyping, variants were again filtered using variant quality score recalibration (VQSR) implemented in *GATK* and only those passing the filter were retained. Given that both studies include samples from both island and mainland areas, and that the number of resulting total sites and variants is comparable across species and populations, the observed differences between island and mainland species are unlikely to result from biased sampling across studies or differences in variant filtering.

We filtered all *vcf* files such that only sites with a minimum Qval of 20 and belonging to autosomal scaffolds/chromosomes were retained. Next, sites were marked as 0-fold, 2-fold, 3-fold and 4-fold degenerate on the basis of the genome annotation of the corresponding reference genome (**Supplementary Table 1**) using the script *NewAnnotateRef.py* by Williamson et al. (2014) available at https://github.com/fabbyrob/science/tree/master/pileup_analyzers. The output file was then converted into bed files for each type of site (0-, 2-, 3-, 4-fold degenerated) using *bedtools,* and a *vcf* file containing these sites was generated. The four resulting *vcf* files were merged and split by chromosome. An additional dataset was produced by excluding CpG-prone sites and weak-to-strong mutations (A-to-G, A-to-C, T-to-G, T-to-C substitutions) following the approach of Kutschera et al. (2020). This dataset was used to assess the effect of *gBGC* by comparison to the dataset including all sites.

### Relatedness and Population Structure

The presence of closely-related individuals, as well as population structure and gene flow between populations, can distort inference of the DFE by mimicking changes in population size, although the effect can be partially absorbed by explicitly modelling a shift in population size, as implemented in Keightley and Eyre-Walker’s (2007) two-epoch model (2e) (see also (Kutschera et al. 2020)). To reduce any effects arising from relatedness or population structure, we first removed related individuals within each species. As a first step, we filtered from the full autosomal dataset sites with linkage-disequilibrium above 60% with *bcftools* (bcftools +prune -l 0.6 -w 50kb -n 5 --AF-tag), and then used plink (v2.0) on the filtered *vcf* file to produce a KING-formatted table of kinship coefficients (option –make-king-table). All individuals with kinship coefficients higher than 0.0442, corresponding to 3^rd^ degree relatives were excluded (Manichaikul et al. 2010).

In order to control for population structure, we additionally performed a PCA for each species using the full set of coding autosomal SNPs and all individuals in *plink* (v1.90b6.21). Many of the individuals appearing as outliers in the first two axes of variation corresponded to those identified as being closely related, providing further arguments for their exclusion from the final dataset (**Supplementary Figure 1**). Moreover, the Italian population of collared flycatchers (*Ficedula albicollis*) and the Spanish population of pied flycatchers (*Ficedula hypoleuca*) were analyzed on their own since they clustered separately from the remaining populations, as has also been previously documented (Burri et al. 2015). Two species of white eyes (*Zosterops borbonicus* and *Z. virens*) were excluded due to significant structure and gene flow between the subpopulations (Bourgeois et al. 2020). Furthermore, as the inference of the site frequency spectrum (SFS) requires sites to be genotyped for all individuals in the sample (i.e. no missing data) we removed individuals with lower quality of sequencing data at the cost of sample size (**Supplementary Table 1**): given that small chromosomes display an increased GC-content and Illumina sequencing biases against them, they are more likely to have missing data (i.e. sites not genotyped for some individuals). Yet, these chromosomes are important for the intended analyses, as they ‘provide’ the high recombination rates, and cannot be discarded. We therefore maximized the number of sites sampled by setting a very conservative threshold for missing data: any individual displaying a fraction of missing data of more than 5% of sites missing in any chromosome was discarded. Moreover, SFS required a minimum of eight chromosomal copies to be sampled, i.e. four individuals with sufficient sequencing coverage (cf. Kutschera et al. 2020). As only exception, the Spanish population of *Ficedula hypoleuca* was included with six chromosomes.

After applying these individual and population filters, the final dataset consisted of 24 populations from 19 species, from which eight were mapped to a reference of the same species, and sixteen to that of the closest relative available in the genus or family. Divergence times between sample and query were within 6.5 mya (**Supplementary Figure 2)** (McTavish et al. 2025). Each population is represented by a median average of 1,604,1053 total and 46,342 segregating sites (see **Supplementary Table 2** for sample sizes of individuals and segregating sites).

### Inference of the DFE

We inferred the folded SFS for each population and each of 28 autosomes separately using the scripts *vcfSummarizer.py* and *bootstrap.py*available at (https://github.com/fabbyrob/science/tree/master/pileup_analyzers). By running these two scripts, all sites with missing data, indels, or multi-nucleotide polymorphisms are discarded. The inferred folded SFS was used to model the DFE of each species/population both per-chromosome and using the full autosomal dataset using *DFEalpha v2.16* (Keightley and Eyre-Walker 2007). This approach requires a set of sites under selection, in which we included the 0-fold, 2-fold and 3-fold degenerated sites, and a set of sites evolving under or close to neutrality, for which we used the set of 4-fold degenerated sites. Two demographic models, one with constant size (1-epoch model) and one with change in population size (2-epoch model) - either an expansion or a contraction - were run for comparison and the best fitting model was selected using a likelihood ratio test. Note, however, that single estimates of long-term *N_e_* are likely not capturing more complex demographic histories including repeated changes in population size with contributions of gene flow. These are likely only controlled in part by the model estimating the DFE from the SFS.

Inference of the best-fitting demographic model differed by chromosome (**Supplementary Figure 3**). This may be due to differences in power but may also suggest that different genomic regions experience demographic variations in different ways, likely due to varying linked selection (see e.g. (Boitard et al. 2022)). Estimates of the DFE can in consequence also be expected to change across chromosomes. Inference of the DFE using the full autosomal dataset was largely dominated by the largest chromosomes, masking different inferences for the smaller chromosomes. Given the variability among chromosomes and our interest in comparing the DFE to recombination rate estimates on a per-chromosome basis, we chose to select the statistically preferred model per chromosome and population rather than imposing a single model to all genomic regions. The fit of the inferred DFE was assessed through the plots of the observed SFS and the ones expected under the best inferred demographic model. These plots are provided in the *zenodo* repository.

While there are alternatives for DFE estimation, like *polyDFE* (Tataru et al. 2017) or *grapes* (Galtier 2016), they turned out to be less suited to the analyses we intended to perform. PolyDFE requires an unfolded SFS which is prone to errors (Tataru et al. 2017) and requires a significant amount of data (around 20Mb), impossible to reach for the smaller chromosomes. Grapes is mainly aimed to estimating the fraction of adaptive substitutions (alpha), and although the DFE is also estimated, the slices it provides (proportions of weakly-and strongly-deleterious mutations) are not directly comparable with the ones provided by DFE-alpha.

### Estimation of the Effective Population Size

Effective populations sizes were obtained from Kutschera et al. (2020) and Leroy et al. (2021), inferred as the harmonic mean of *N_e_* estimates derived from the sequential Markovian coalescent (McKenna et al. 2010; Schiffels and Durbin 2014) weighted by the length of each time period. For details see Supplementary Text S1 in Kutschera et al. (2020) and Method detail in Leroy et al. (2021). According to the neutral theory, long-term coalescent *N_e_* should be similarly reflected by *θ*_π_ *= 4N_e_µ.* In agreement with this prediction, estimates from the sequential Markovian coalescent covaried strongly with estimates of synonymous nucleotide diversity (π_s_) as in Leroy et al (2021). The estimates were obtained from pairs of genomic sequences reconstructed for each individual, applying filters for depth of coverage, and removing sites with missing data (**Supplementary Figure 4**). Note that both estimates of *N_e_* comprise the entire population history likely including a mainland phase prior to island colonization in several cases. Since the DFE was equally shaped during the entire history of the population, including the mainland phase, these long-term estimates of *N_e_*, however, seem an appropriate choice.

Besides these coalescent-based estimates, we used geographical range (Island vs. Mainland) as a proxy to *N_e_* (see also discussion). This assumption is based on the expectation that species occupying mainland territories have larger population sizes than those living on the smaller territories located in islands (**Supplementary Figure 5**). Moreover, this link is further supported by the documented observation, in agreement with the predictions of neutral theory, that populations with large *N_e_* should be more genetically diverse than those with reduced effective population sizes (Kutschera et al. 2020; Leroy et al. 2021) (**Supplementary Figure 4)**. Life history traits influence *N_e_* and evolutionary rates, and might also affect the DFE (Romiguier et al. 2014). Since all species under considerations here are passerines with comparable life histories, we abstained from including this in the statistical models to avoid overparameterization.

### Recombination Rate Estimation

The average recombination rate (*r)* per chromosome was approximated in centimorgans per mega base pair [cM/Mb] as follows: we extracted estimates of the per-base population recombination rate ρ[1/bp] obtained by Vijay et al. (2016) in bins of 50 kb windows for 3 populations of hooded and carrion crows (*Corvus (corone) ssp.*). We then calculated, for each population, the median across the 50 kb windows to obtain per-chromosome estimates. Last, we averaged the estimates across the three populations. We then estimated genetic variation along each chromosome and converted the population recombination rate estimates ρ[1/bp] to recombination estimates in centimorgan using the relationship ρ */* π *= 4N_e_r* / *4Neµ = r / µ.* Multiplication with the mutation rate and scaling to cM (10^2^) and Mb (10^6^) then yields r[cM/Mb]. We chose to use a single mutation rate of 4.6x10^-9^ mutations/site/generation as estimated by Smeds et al. (2016) for the collared flycatcher. The resulting per-chromosome estimates range between 2.46 and 11.5 cM/Mb (mean weighted by chromosome length: 3.57) adding up to a total genetic map length of 3,500 cM which is comparable to previous, pedigree-based estimates of passerine birds (Backström et al. 2010; Kawakami et al. 2014). The choice of a single mutation rate for all species seems justified, as the estimate for flycatcher is highly comparable to that of zebra finch (5.0 × 10-9 mutations/site/generation) (Prentout et al. 2025). Flycatchers (Muscicapidae) and zebra finches (Estrildidae) diverged ∼21 mya ago, which approximates the range of 28 mya of divergence for our study organisms (**Supplementary Figure 2**) (McTavish et al. 2025).

Central to our study is the assumption that the karyotype of passerine species considered here is stable enough to reflect comparable recombination rates across syntenic chromosomes (Dutoit et al. 2017; Vijay et al. 2017). To test the assumption that a single set of recombination rates is representative for the species involved in the analyses, we compared chromosome-level recombination rates of hooded crow (Vijay et al. 2016) with those of zebra finch (*Taeniopygia guttata*) (two estimates from Singhal et al. (2015) and Stapley et al. (2010)), collared flycatcher (*Ficedula albicollis*) (Chase et al. 2024), and Eurasian blackcap (*Sylvia atricapilla*) (Bascón-Cardozo et al. 2024). Analogous to the mutation rate estimates for flycatcher and zebra finch described above, these species represent approximately 23 million years of avian evolution (divergence time Sylvidae – Estrildidae) and thus encompass most of the divergence among the species included in this study. As the estimates of zebra finch in Singhal et al (2015) and the ones of the Eurasian blackcap were provided as estimates of the population recombination rate (ρ / *bp*), we used the values of genetic diversity provided in the respective studies (genomic estimate of 0.082 for zebra finch; per chromosome estimates in Eurasian blackcap as in Table S3 of (Bascón-Cardozo et al. 2024)), and the avian mutation rate as in Smeds et al. (2016)(4.6x10^-9^ mutations/site/generation), to convert them into cM/Mb using the same approach as for hooded crow outlined above.

Chromosome-level recombination rates were significantly correlated across all comparisons (median R^2^=0.668, median p-value = 1.838787^-07^, **Supplementary Figure 6**) supporting that, at chromosome-level, recombination rates are comparable across passerine birds. A single recombination estimate, here from crows, appears thus suitable for comparative analysis, although it will introduce noise which may conceal the effect of recombination rate on the DFE. Moreover, chromosome length strongly correlates with recombination rate and is highly conserved across avian species (Ellegren 2010, this study; Bascón-Cardozo et al. 2024). We therefore also used chromosome length as a proxy for recombination rate.

In order to ensure that the chromosomes correspond to the names assigned in the different references used here (**Supplementary Table 1**), we verified synteny based on the corresponding annotations using custom scripts. First, all annotated genes for each chromosome were text-mined from annotation files for hooded crow, zebra finch, collared flycatcher, great tit and chaffinch. The gene lists were then compared for each chromosome between each possible pair of references. The annotations were highly consistent and synteny could be verified, with few exceptions, for all references. Most mismatches occurred between the reference for hooded crow and the other references, mainly due to crow scaffolds belonging to a) chromosomes that were not present in the crow assembly version GCF_000738735.1, to which the *vcf* files utilized here correspond, being assigned to other chromosomes (e.g. 25, LGE22) or to b) small chromosomes (e.g. 23, 27). The identified mismatches were corrected and the scaffolds included in the correct chromosomes, if they were confirmed to be syntenic across all the other analyzed references; or excluded, if the chromosomes were not present in the hooded crow assembly. A few mismatches from the used zebra finch reference (GCF_000151805.1) have been already corrected and verified as syntenic in a more recent annotation (GCF_003957565.2).

### Statistical Analyses

We used a mixed linear model framework to test the influence of recombination (*r*), effective population size (*N_e_*) and GC-biased gene conversion (*gBGC*) on the proportion of mildly deleterious alleles (*N_e_s* ∈ (-1;0]). Recombination and effective population size were approximated by two alternative measures (*r*: recombination rate [cM/Mb] or chromosome length (see above); *N_e_*: species restricted to island yes/no or log(*N_e_*)). The main results were largely unaffected by the choice of measure. In the main text, we present the results for recombination rate [cM] and island yes/no. Results for the alternative measures can be found in **Supplementary Table 3.** The effect of *gBGC* was assessed as categorical variable by comparing the estimate of DFE using all polymorphisms (with *gBGC*) or the subset including only weak-to-weak (A-T), and strong-to-strong mutations (C-G) (without *gBGC*).

We fitted a mixed effect model including all three parameters and a choice of meaningful interactions as fixed effects using the lmer function of the R package lme4 (Bates et al. 2015). We were interested whether the effect of recombination on the DFE depended on *N_e_* or *gBGC* (pairwise interactions: *r*N_e_, r*gBGC*), as well as whether the effect of *gBGC* differed by *N_e_* (pairwise interaction *N_e_ *gBGC*). In addition, to understand whether the relative rates of the two recombination-mediated, opposing processes of purging and *gBGC* differed by *N_e_*, we also investigated the interaction of all three factors (three-way interaction: *r*N_e_*gBGC*). We further incorporated chromosome (chr) and population identity (pop) as random effects including variation in slope for the latter. More than a single population was available for a subset of five species, which led us to also include species (spec) as a nested random effect (population nested in species). Model choice was then performed largely following (Zuur et al. 2009): we started with a model containing the full set of random factors and all relevant fixed factors and their interactions: *DFE ∼ r + Ne + gBGC + r*Ne + r*gBGC + Ne*gBGC + r*Ne*gBGC + (1|chr) + (1+r|pop/spec).* We then selected the best combination of random parameters using restricted maximum likelihood estimation (REML) minimizing the Akaike Information Criterion (AIC) which eliminated the nested population/species component and reduced the random effects to *(1+r*|*pop)+(1|chr).* Next, we formulated a set of 10 models exploring the influence of *r*, *N_e_*, *gBGC* and their interactions with inclusion of the null model (see **Supplementary Table 3**) and selected the model minimizing AIC using full maximum likelihood which provides information about the goodness of fit of the respective model to the observed data and penalizes overly complex models (Johnson and Omland 2004). Models differing by less than two units of AIC were considered to be supported. For parameter estimation, we then re-ran the best model using REML estimation and conducted an analysis of variance as implemented in the R package car (Fox and Weisberg 2019). Finally, we assessed normality of residuals by visual inspection and model performance by plotting observed to predicted values (**Supplementary Figure 7**).

In order to assess the impact of phylogenetic dependency, we built a mitochondrial phylogeny for all 19 species using part of the sequences included in Leroy et al (2021), to which we added missing sequences for the two crow species (NC062298.1, NC051471.1). We inferred a maximum likelihood (ML) tree using *IQtree* (v2.1.3) (Minh et al. 2020) under a GTR+GAMMA+I substitution model to account for base content and rate heterogeneity across sites (**Supplementary Figure 8**). We then fitted Bayesian mixed models using the MCMCglmm package in R (Hadfield 2010). We modeled the DFE as a function of recombination rate (parametrized as *r* [cM/Mb]) and *N_e_* (parameterized as island yes/no), *gBGC* (yes/no) and their interactions, while including a phylogenetic random effect whose covariance structure was specified by the inverse of the phylogeny-derived relatedness matrix. Following the mixed model above, chromosome was added as additional random factor. We focused model comparison on the null model and the two best models as inferred by the mixed model described above (three-way interaction, all two-way interactions). Models, including only one population per species (see **Supplementary Table 2**) were run under Gaussian error structure using weakly informative priors (G: *V* = 1, ν = 0.002; R: *V* = 1, ν = 0.002) with 150,000 iterations, a 50,000-iteration burn-in, and thinning interval of 50 for which no evidence of autocorrelation remained. Model choice was based on the Deviance Information Criterion (DIC) and visual inspection of posterior distributions. In this Bayesian framework, effects with posterior distributions that exclude zero correspond to high posterior probability of an effect, analogous to low *p*-values in frequentist analyses, while broadly overlapping distributions indicate weak or no support.

## Results

We calculated the DFE for each of the 24 populations from 19 different species based on segregating sites in coding sequence. The DFE was calculated separately for each of 28 autosomes by contrasting the site frequency spectrum (SFS) of neutral mutations at 4-fold degenerate sites with the frequency of non-synonymous mutations subject to selection controlling for demographic perturbation where necessary (see methods). Recombination rate, approximated in cM/Mb, covaried negatively with chromosome length and (*gBGC*-impacted) GC content (log-log correlations: R^2^=0.77 and 0.81, respectively; both p<0.001) (**Figure 2**).

In order to establish a baseline without the influence of *gBGC*, we first performed the analyses on the SFS after excluding CpG-prone sites and weak-to-strong mutations [A,T]->[C,G]. The median, genome-wide proportion of mildly deleterious mutations differed substantially between populations (min/median/max: 0.09/0.18/0.25), as well as between species (0.09/0.19/0.25). This variation in mutation load appears to be well explained by population size, regardless of whether it was approximated by coalescence-based estimates of *N_e_*or by species distribution (island vs. mainland) (**Figure 3A, Supplementary Figure 5**). Species restricted to islands had a median average *N_e_* of approx. 90,000, while for mainland species this value was approx. 160,000. This 2-fold difference in *N_e_* was reflected by a median increase in the mutation load of approximately 1.6-fold. Mutation load was not only dependent on *N_e_* of the species, but also varied along the genomes of nearly all studied populations as a function of recombination rate: low-recombining chromosomes carried substantially more load than high-recombining genomes for both island and mainland populations (**Figure 3A**).

**Figure 3.**
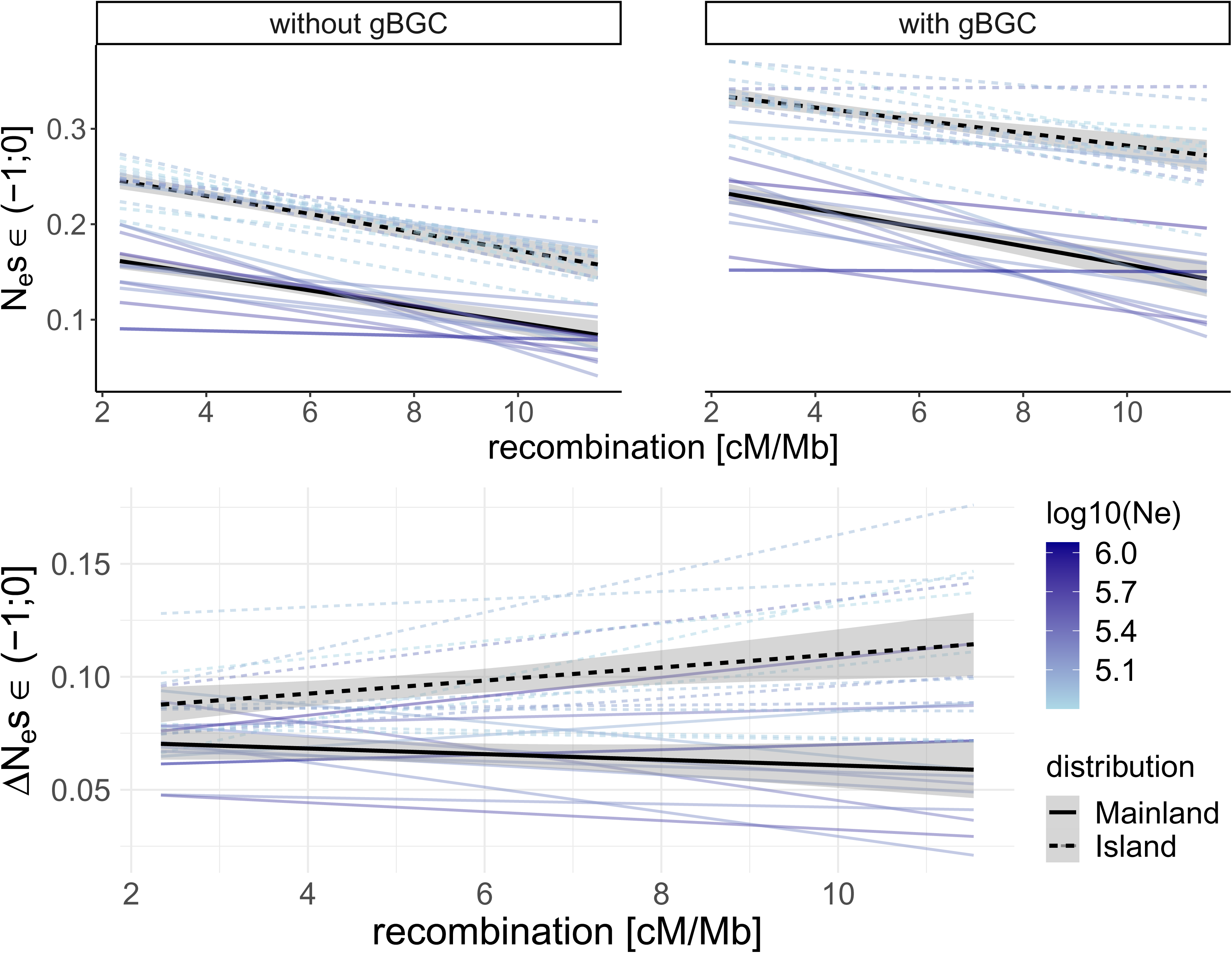
Relationship between recombination rate and mutation load estimated as the fraction of mildly deleterious mutations with N_e_s ∈ [-1;0). Correlation between recombination rate and mutation load A) excluding and B) including mutations subject to *gBGC* (right). Each regression line represents a population sample of a species differing in effective population sizes (*N_e_*) as is indicated by the blue colour gradient on a logarithmic scale. Dashed and solid lines illustrate island and mainland populations, respectively. C) as above, but the y-axis displays the difference between estimates of mutation load with and without mutations subject to *gBGC*. Note that the slopes differ between island and mainland populations indicating that relative rates of the two recombination-mediated processes, *gBGC* and purging efficacy, may interact with the effective population size of a species.

*gBGC* increased the mutation load by approximately 1.5-fold on median average (**Figure 3B**). At first glance, the patterns of mutation load in relation to recombination rate and species-level effective population size (*N_e_*) appear similar whether or not biased gene conversion is considered (**Figure 3A** vs. **Figure 3B**). To formally evaluate the effects of population size, recombination, *gBGC*, and their interactions on mutation load, we employed a linear mixed-effects model (**Supplementary Table 3**). Chromosome and population identity were included as random effects, as they accounted for a significant proportion of additional variance. The best-fitting model (**Table 1, Supplementary Figure 7**) indicates a strong influence of all three factors, summarized as follows: (i) mutation load was substantially higher in small populations, (ii) was further increased for mutations affected by *gBGC*, and (iii) was markedly reduced in genomic regions with high recombination rates. The model also predicts that the contribution of *gBGC* to deleterious variation is greater in small populations (i.e., a significant *N_e_ x gBGC* interaction), with island populations showing a 9.6% higher proportion of deleterious mutations compared to a 6.4% increase in mainland populations. Nonetheless, the relative increase due to *gBGC* remained consistent across population sizes, with island and mainland populations exhibiting a similar fold-increase in deleterious mutations (1.44-fold vs. 1.46-fold, respectively).

**Table 1:** Summary of the four best statistical models and the null model exploring the relationship between the mutation load quantified as the proportion of mildly deleterious alleles (DFE) with recombination rate (r), effective population size (Ne) and GC-biased gene conversion (*gBGC*). *r* was here approximated by chromosome length, N_e_ was here approximated by whether a population was restricted to an island (yes/no), *gBGC* was estimated for all sites (with *gBGC*) and for a subset of sites excluding weak-to-strong mutations (without gBCG). For other proxies of r and N_e_ and the full set of models with parameter estimates see **Supplementary Table 3.** Model selection is based on Akaike’s Information criterion (AIC) with models by ΔAIC_c_<2 are considered supported (best model in bold). In addition, thresholds of Type 1 error probabilities of p<0.001 (***), p<0.01 (**), p<0.05 (*) and p>0.05 (n.s) are indicated for all fixed effects and their interactions included in a specific model.

| Model | $r$<br>[cM/Mb] | $N_e$<br>[yes/no] | $gBGC$<br>[with/without] | $N_e * gBGC$ | $r * N_e$ | $r * gBGC$ | $r * N_e * gBGC$ | k | AIC | $\Delta AIC$ |
| --- | --- | --- | --- | --- | --- | --- | --- | --- | --- | --- |
| **10** | *** | *** | *** | *** | n.s. | n.s. | * | **13** | **-4479.5754** | **0** |
| 9 | *** | *** | *** | *** | n.s. | n.s. | - | 12 | -4477.2763 | 2.299 |
| 8 | *** | *** | *** | *** | - | - | - | 10 | -4476.8571 | 2.718 |
| 7 | *** | *** | *** | - | - | - | - | 11 | -4440.2616 | 39.31 |
| NULL | - | - | - | - | - | - | - | 6 | -3555.9041 | 923.67<br>1 |

The data further allowed us to assess the relative rates by which the two opposing, recombination-dependent processes of purging and *gBGC* affect mutation load. If both processes took place with the same intensity, then a doubling in the increase of deleterious mutations by *gBGC* should be fully offset by a proportional increase in purging efficacy. The relative contribution of both processes may further depend on the species’ *N_e_* (**Figure 1**). We found that the purging efficacy of recombination on mutation load was reduced in small populations for the mutational spectrum affected by *gBGC*. In other words, small populations suffer from a disproportionate introduction of deleterious [A,T]->[C,G] mutations that is not fully offset by purging efficacy in regions of high recombination (**Figure 3C**). This effect (three-way interaction *r x N_e_ x gBGC*) was weak but statistically supported when using a binary measure of population size (island/mainland) (**Table 1**, ΔAIC to 2^nd^ best model =2.229), but not when parametrizing over a continuum of coalescent based estimates of *N_e_* (**Supplementary Table 3**). The three-way interaction using the island/mainland contrast still had support when controlling for phylogenetic dependence using a dataset limited to the 19 species in a Bayesian approach (ΔAIC to 2^nd^ best model =1.492, p-value analogue=0.059, see **Supplementary Table 3**).

## Discussion

In this study, we estimated the distribution of fitness effects for 24 populations from 19 avian species including representatives from across the major songbird clades (Corvoidea, Muscicapoidea, Passeroidea, Sylvoidea) (Jønsson and Fjeldså 2006). Oscine songbirds are the most speciose avian clade arising broadly around the Cretaceous-Palaeogene boundary some 66 million years ago (Sibley and Ahlquist 1990; Barker et al. 2004; Jarvis et al. 2014). The study thus claims some generality at the higher end of evolutionary timescales compared to previous comparative work in the area (within species: (Eyre-Walker et al. 2006; Slotte et al. 2010; Deinum et al. 2015; Castellano et al. 2018), genera: (Kutschera et al. 2020), families: (Castellano et al. 2019), and beyond (Chen et al. 2017; Leroy et al. 2021; Chen et al. 2022)). With the exception of few closely related species, the chosen species in this study are overall phylogenetically distant enough as not to share ancestral polymorphism (isolation > 9 Ne generations Hudson et al. 2002; Mugal et al. 2020). As some evolutionary dependency may remain, we additionally fitted a model taking phylogenetic contrasts into account.

Focusing on the mildly deleterious proportion of the DFE, we tested the effects of *N_e_*, recombination rate, *gBGC*, and their interaction, on mutation load. The effective population size, as parameterized by species distribution (island/mainland) or coalescent-based estimates, was a strong predictor. Coalescent effective population sizes of island populations were approximately half of the mainland populations. In accordance with the central tenet of the nearly-neutral theory and previous studies, mutation load was significantly higher in small effective populations (Ohta 1992; Charlesworth 2009; Agrawal and Whitlock 2012; Kutschera et al. 2020; Leroy et al. 2021). Our model predicts this difference to translate into an increase in the genome-wide mutation load of approximately 1.6-fold on islands (cf. (Castellano et al. 2018) for a similar estimate considering local genomic N_e_ variation within the *Drosophila* genome). While the difference in mutation load on islands is mediated by small population size, a rather discrete change between mainland and island populations may also suggest additional factors – such as bottlenecks or different positions in the fitness landscape (see below) – that are not fully accounted for by the coalescent-based *N_e_* estimates (**Supplementary Figure 4**). Moreover, single estimates of long-term *N_e_* are likely not capturing more complex demographic histories including repeated changes in population size with contributions of gene flow. These are likely only controlled in part by the model estimating the DFE from the SFS.

Despite these limitations, these results add to the evidence that species living on islands accumulate mildly deleterious mutations more readily than their mainland counterparts (Johnson and Seger 2001; Woolfit and Bromham 2005; Kutschera et al. 2020),making them more vulnerable to extinction in the long term (Frankham 1998). Our results further suggest that–mutation load is unevenly distributed across the genome. In accordance with relatively few previous studies touching upon the subject (Gossmann et al. 2011; Leroy et al. 2021; Chase et al. 2024), we find a clear reduction of mutation load with an increase in recombination. In our model, the consistent ∼ 5-fold range of recombination rate across avian chromosomes (Backström et al. 2010; Kawakami et al. 2014; Bascón-Cardozo et al. 2024, this study) translates to a ∼1.7-fold change in mutation load. The order of this change is comparable to the effect of *N_e_*-change between island and mainland. Importantly, however, the effect of recombination was independent of *N_e_* (no statistical interaction *N_e_ x r*) suggesting the same relative purging efficiency between chromosomes regardless of population size. This result is in conflict with conclusions from Leroy et al. (2021). While they also find reduced mutation load in genomic regions of high recombination, the beneficial effect of recombination was stronger in island populations. This disconnect may be explained by a combination of factors like using GC content at the third position of the codon (GC3) as proxy for recombination, a different set of species and presence of population structure in several of their datasets that can influence the DFE (Kutschera et al. 2020) and which we removed for this study (**Supplementary Figure 1**).

In addition to *N_e_* and intragenomic heterogeneity of recombination rate, the quality of mutations also affected mutation load. When including mutations subject to *gBGC* the genome-wide proportion of mildly deleterious variation increased by ∼1.5 fold, both in island and mainland populations. An increase of mutation load in this fraction of mutations is expected, as they are replenished through the segregational bias mimicking the effect of positive selection (Berglund et al. 2009). Yet its interaction with recombination and overall population size remains largely unclear. Since *gBGC* relies on double-strand breaks associated with increased levels of recombination, *gBGC* mediated mutation load is expected to be more pronounced in regions of high recombination (Duret and Galtier 2009; Kostka et al. 2012; Pouyet et al. 2018). We thus expected to see relatively more deleterious mutations in regions of high recombination with *gBGC* than in those with lower recombination rates (flatter slope for DFE with *gBGC*, **Figure 1**). We observed the effect only in island populations, while the effect may be too slight to pick up in the mainland with an overall increased efficacy of purifying selection. Alternatively, the relative increase in *gBGC*-contributed mutation load in small island populations, may reflect the reduced overall intensity of *gBGC*. Genes experiencing high levels of *gBGC* in regions of high recombination tend to incur a fitness disadvantage (Joseph 2024), but can still accumulate deleterious AT-GC mutations in large populations where the *gBGC* pressure is high. In small populations, reduced *gBGC* pressure might facilitate segregation of GC→AT back mutations in regions of high recombination. The outcome depends on the relative effect of recombination on *gBGC* and selection efficacy.

Returning to the hypotheses of the introduction, we conclude that mutation load 1) is elevated in (island) populations with low *N_e_*, 2) is reduced in highly-recombining regions, and 3) is further increased for the mutational spectrum subject to *gBGC*. All three effects are of comparable magnitude across the relevant biological range of variation in our sample. Relative rates of the two recombination-mediated, opposing processes of purging and *gBGC* differ in populations with low *N_e_*. Only few studies have hitherto explored these interactions (Jiang et al. 2025). In our view, they deserve more empirical and theoretical attention, as the fundamentally affect the overall magnitude and genomic distribution of mutation load segregating in natural populations.

In addition, non-negligible variance for the species random effect is consistent with additional components affecting a population’s’ DFE, such as its current position in the fitness landscape (Lanfear et al. 2014). For species at or close to the fitness optimum, most mutations can be expected to have deleterious effects, whereas in species further removed from their fitness optimum, a subset of mutations may have beneficial effects. This may also impact our proxy for *N_e_*. While geographic restriction has been suggested as a valuable proxy for the effective population sizes (Kutschera et al. 2020; Leroy et al. 2021), island or mainland populations may qualitatively differ in their position in the fitness landscape and thereby in the mutation load they experience.

Overall, this study highlights that assuming a homogeneous DFE across the genome is a substantial oversimplification of the complex interplay among evolutionary forces shaping genomic diversity. As a consequence, genetic risk factors are non-homogeneously distributed across the genome (Kardos et al. 2021). Emerging medical, population and conservation genomic studies screening for deleterious mutations and their fitness effects (Bertorelle et al. 2022; Hoffman et al. 2024; Chen et al. 2025) may benefit from taking the interaction between recombination, *gBGC* and *N_e_*into account.

## Data Availability

Whole genome resequencing data are available through the National Center of Biotechnology Information (NCBI). Accession numbers are provided in **Supplementary Table 1**. Analysis scripts, processed *vcf* files and plots of the SFS for individual chromosomes for each population are available at the zenodo repository (doi: 10.5281/zenodo.17822666).

## Acknowledgements

We thank Benoit Nabholz and Verena Kutschera who provided access to raw data and *vcf* files. We also acknowledge valuable feed-back from Dirk Metzler on the Bayesian statistics.

## Author Contributions

JBWW and FB conceived of the study. FB conducted all bioinformatic analyses, JBWW conducted the statistical analyses. JBWW and FB wrote the manuscript.

