## Supplementary Text and Figures for "Determinants of mutation load in birds"

**INDEX**

Supplementary Figures	page 4

Supplementary Tables	page 17

**Supplementary Figures**

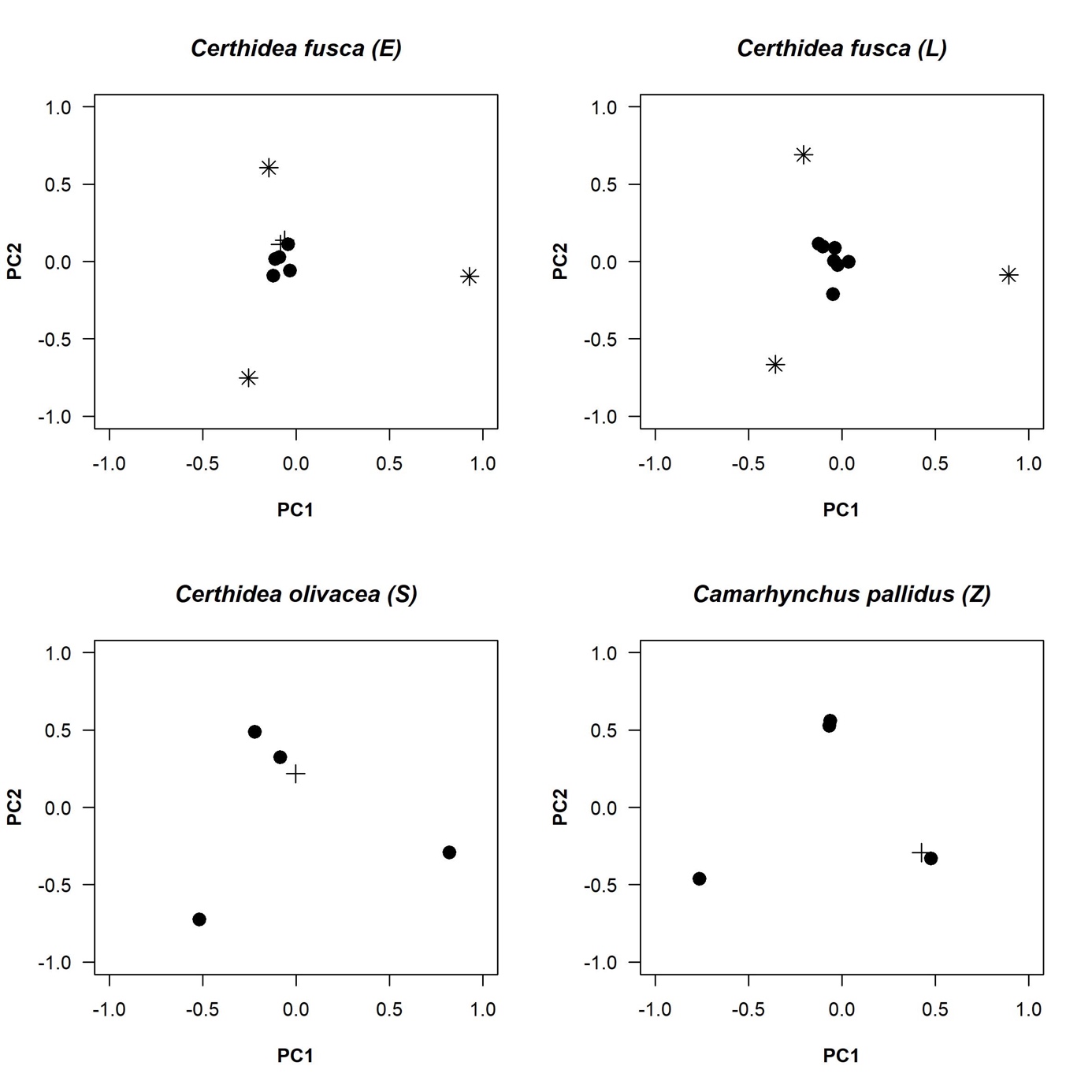

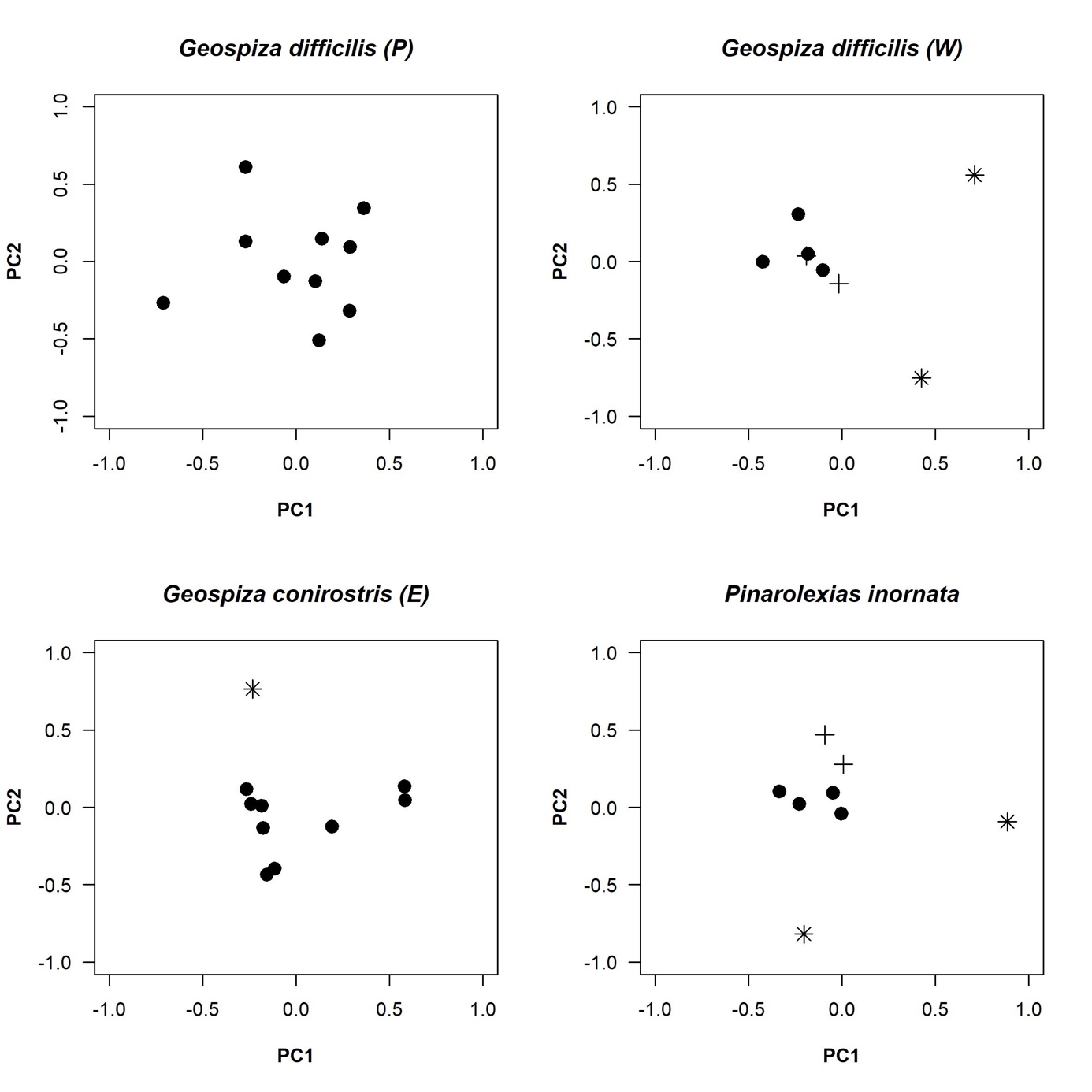

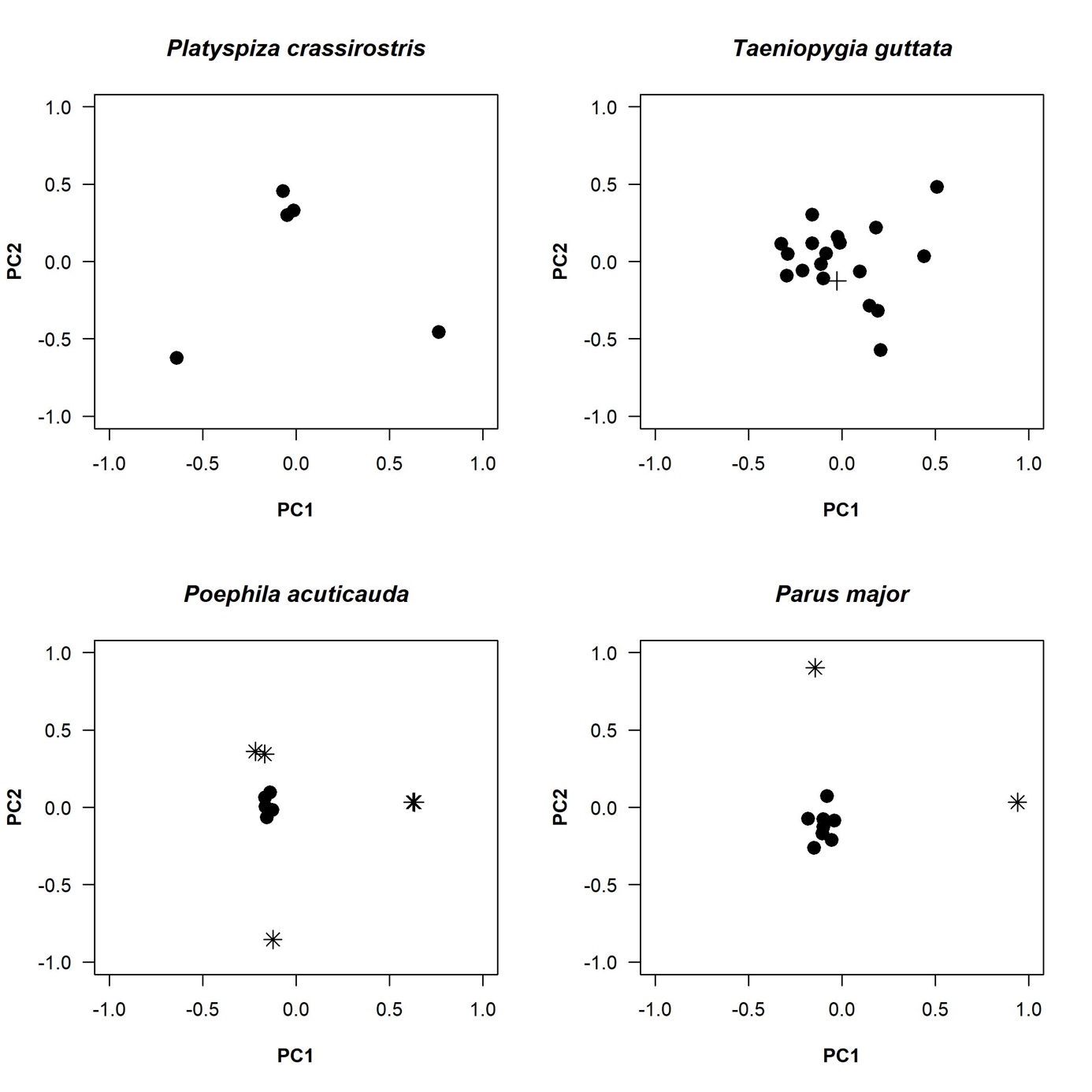

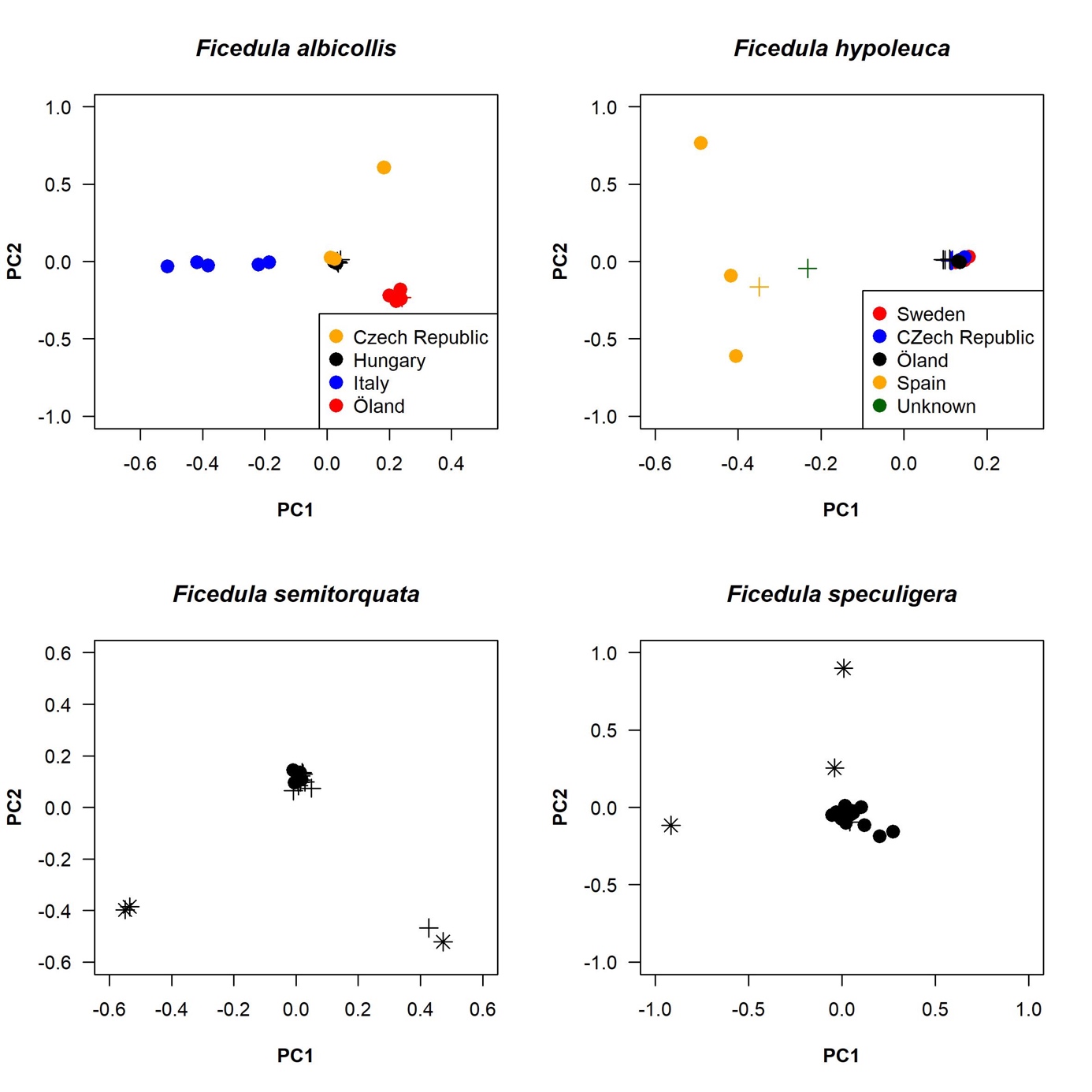

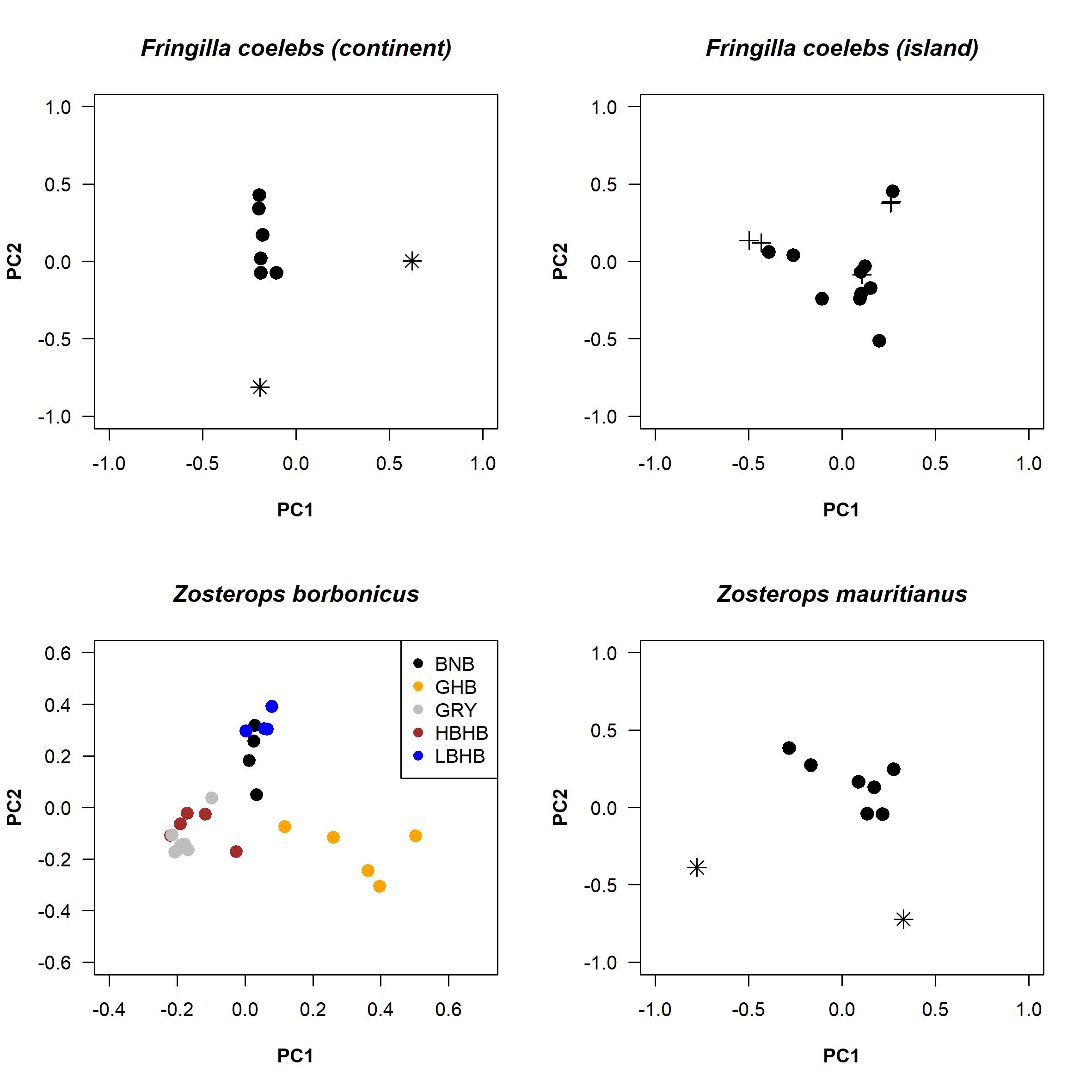

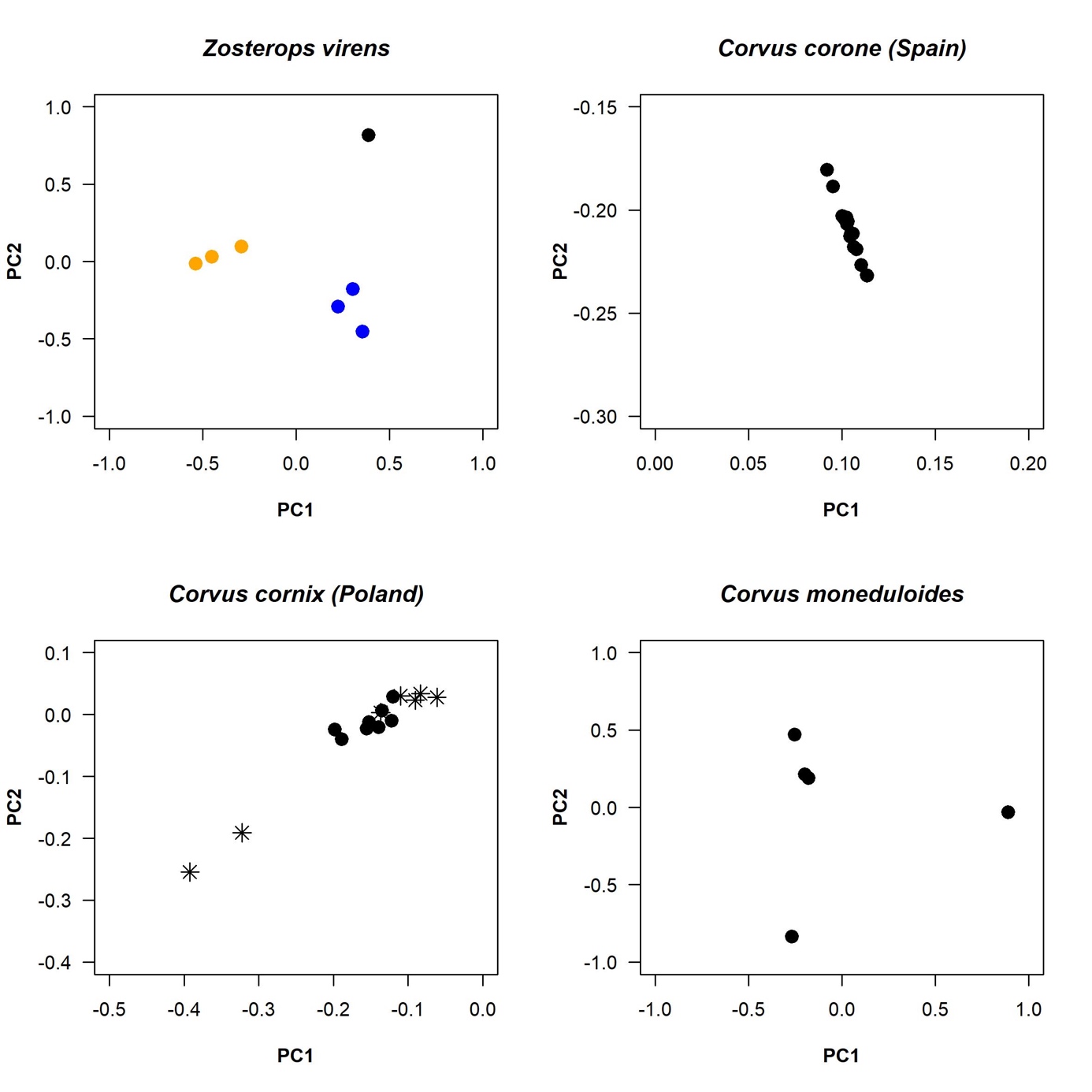

**Supplementary Figure 1 |** **Population structure for all populations and species.** Principle component analysis was conducted to assess population structure. Species with evidence for population stratification were either dropped, or subsampled to a population with at least 4 individuals (exception Spanish Ficedula flycatchers with n=3). Samples that have been removed from the final analysis on the basis of PCA analysis are shown as an asterisk, those removed because of missing data are indicated with a cross. Colored points refer to geographic population samples.

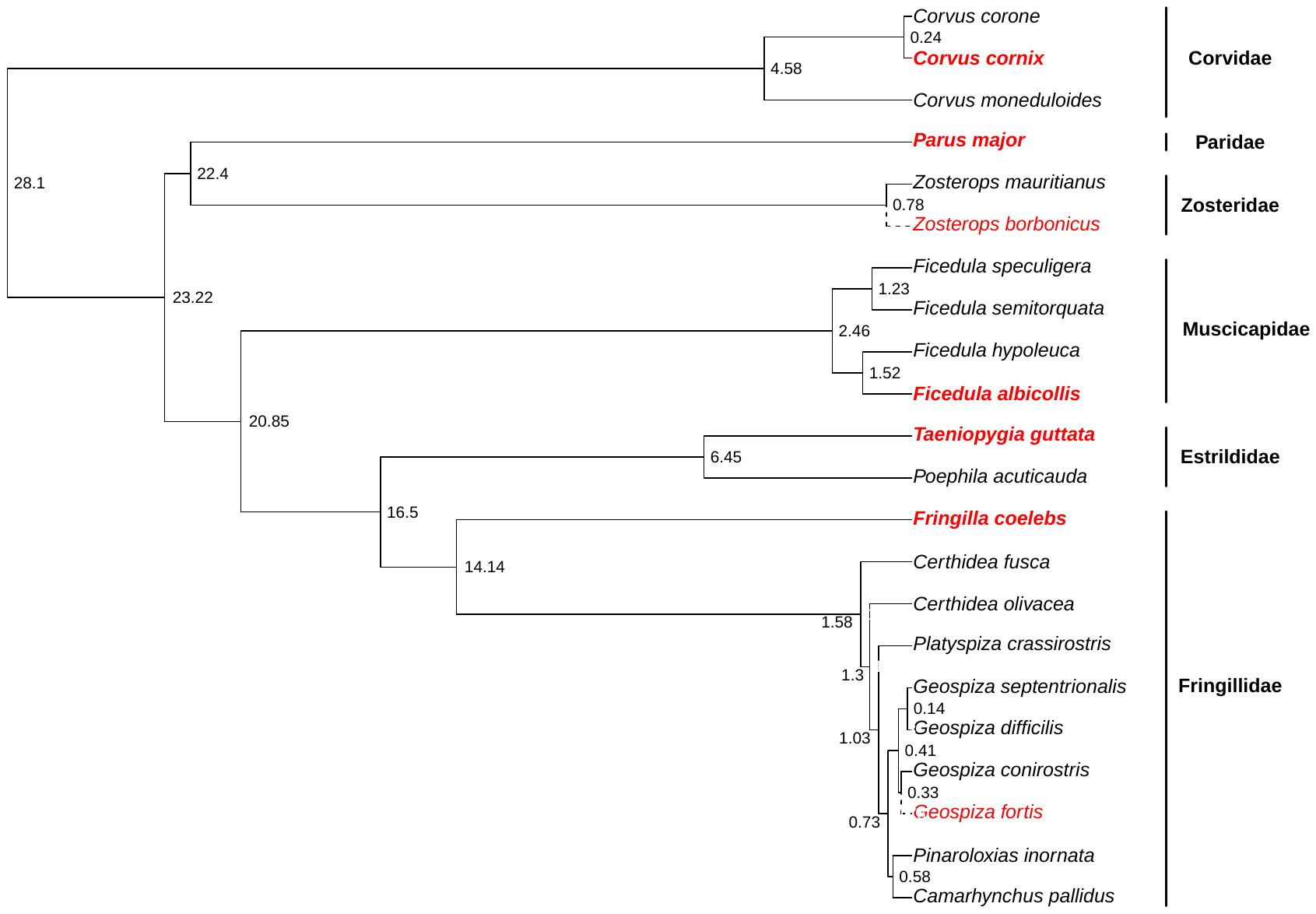

**Supplementary Figure 2 |** Pruned avian phylogeny from (McTavish et al. 2025) showing all species used for this study. Species for which no reference genomes were available (black) were mapped to the closest available references (red). Species with dotted tips were merely used as reference genomes and have not been included in the evolutionary analyses of this study. Contrary to McTasvish et al. 2025, we treat Corvus corone and C. cornix as a single species.

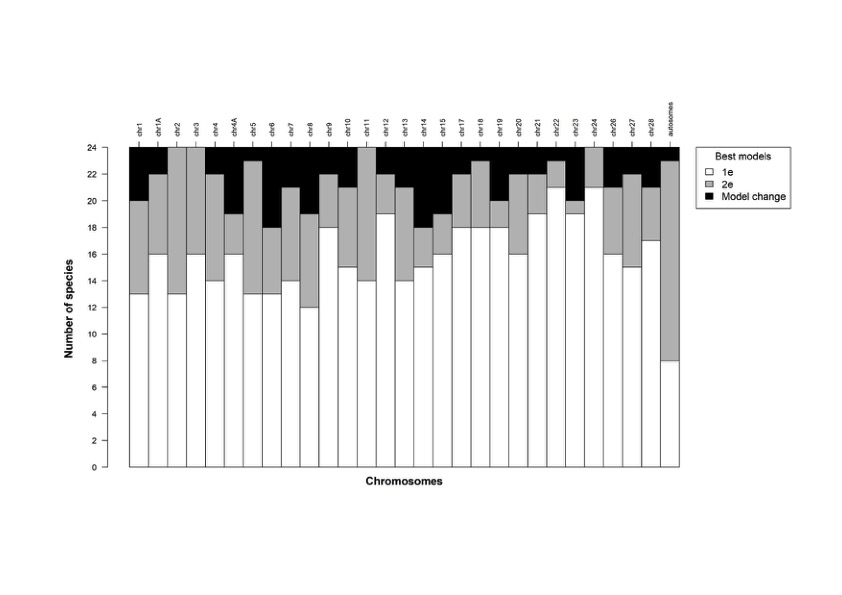
 **Supplementary Figure 3 |** **Chromosome-specific inference of demographic perturbation on the site frequency spectrum.** The presence of population structure (excluded here), demographic perturbation or linked selection can distort inference of the distribution of fitness effects. Keightley and Eyre-Walker’s (2007) 2-epoch (2e) model at least partly absorbs this effect. Inference of the best fitting model differed by chromosome and population. For the majority of populations, the 1-epoch model (1e) was preferred for all chromosomes, both when using all sites or excluding sites prone to GC-biased-gene conversion. In few cases, the best model differed between these two data sets (Model change). When all autosomes were considered together, the 2-epoch model was preferred in most species. For the final analyses we chose the model that was best supported for each chromosome and population.

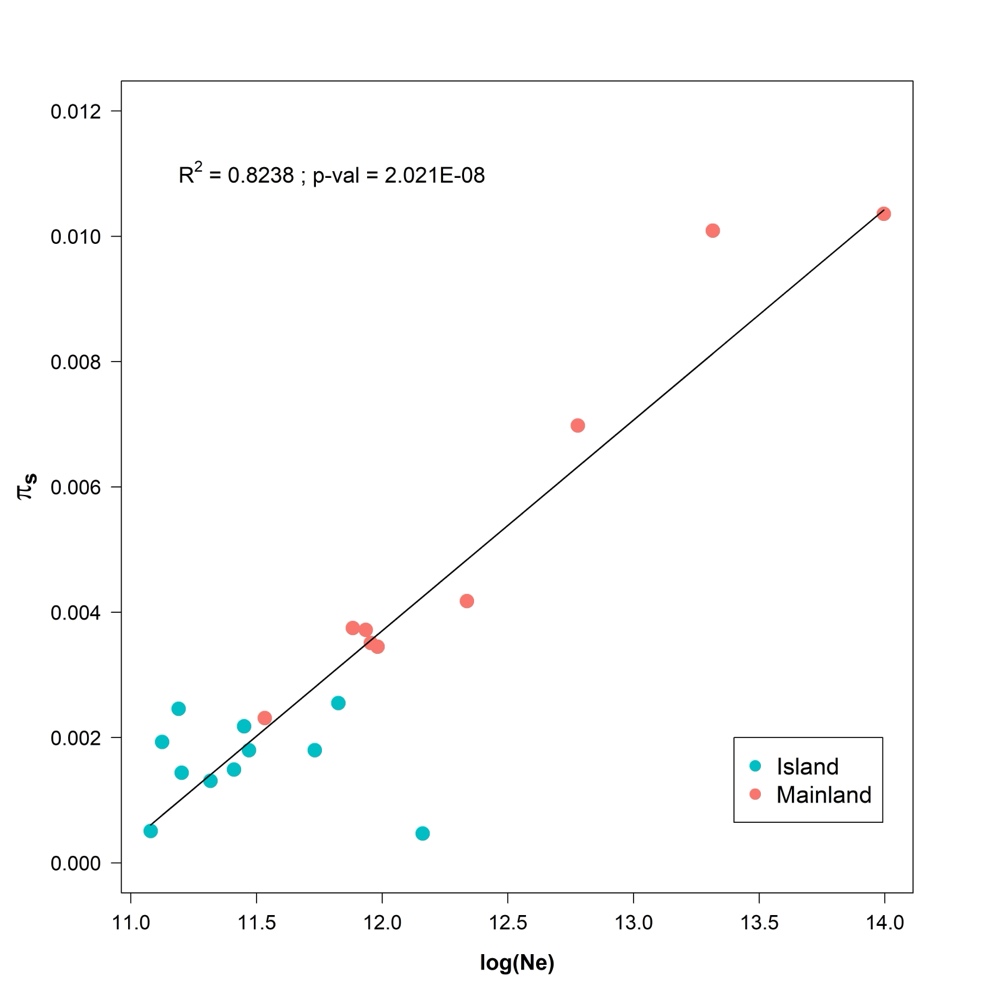

**Supplementary Figure 4 | Relationship between effective population size (log *N_e_*) and synonymous genetic diversity (*π_s_*), for populations distributed on the mainland (red) and island (blue).** Populations occupying the larger mainland territories have higher genetic diversity than those inhabiting islands, as predicted by the neutral theory.

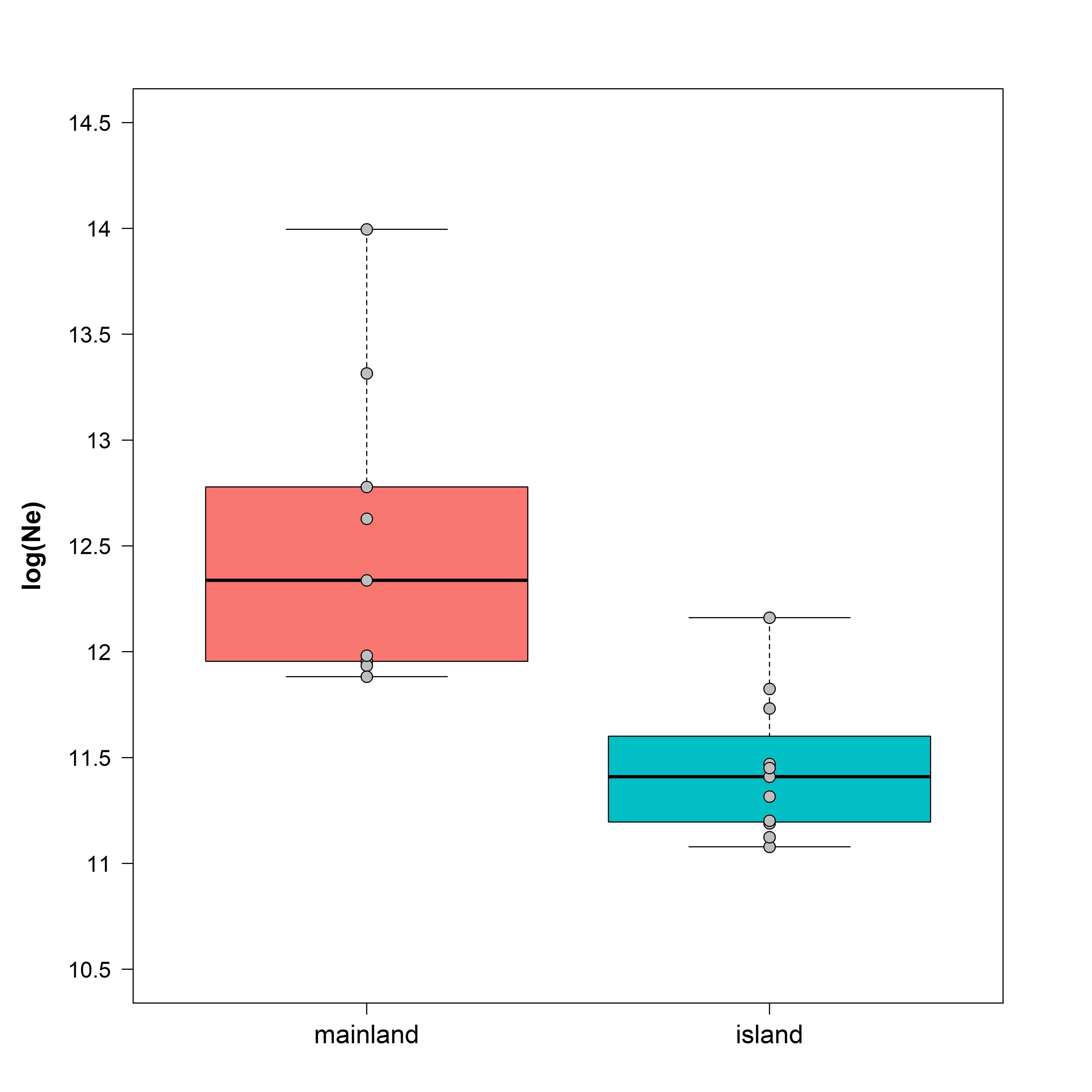

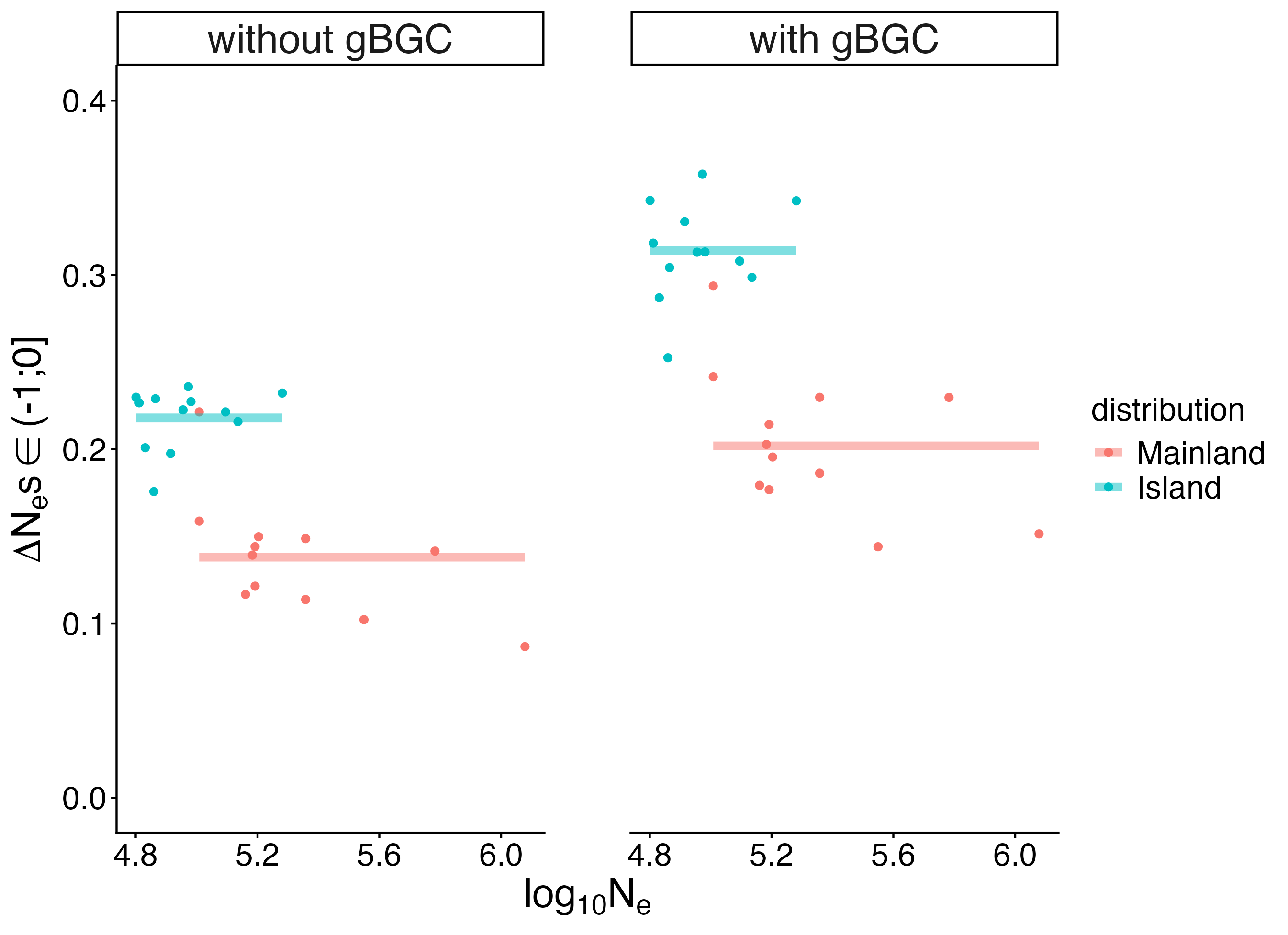

**Supplementary Figure 5 |**

**Left: Comparison of effective population sizes between populations with mainland and island distributions.** Gray dots represent the actual values estimated for the populations, and thick horizontal lines indicate the median value.

**Right: Relationship betw**e**en the effective population size (*N_e_*) and the proportion of mildly deleterious mutations (*N_e_s* ∈ (-1;0]; DFE).** The left panel shows estimates excluding CpG-prone sites and weak-to-strong mutations [A,T]->[C,G] subject to biased gene conversion, the right panel includes all mutations. Colors indicate whether a species is restricted to an island (red) or has a wider mainland distribution (blue).

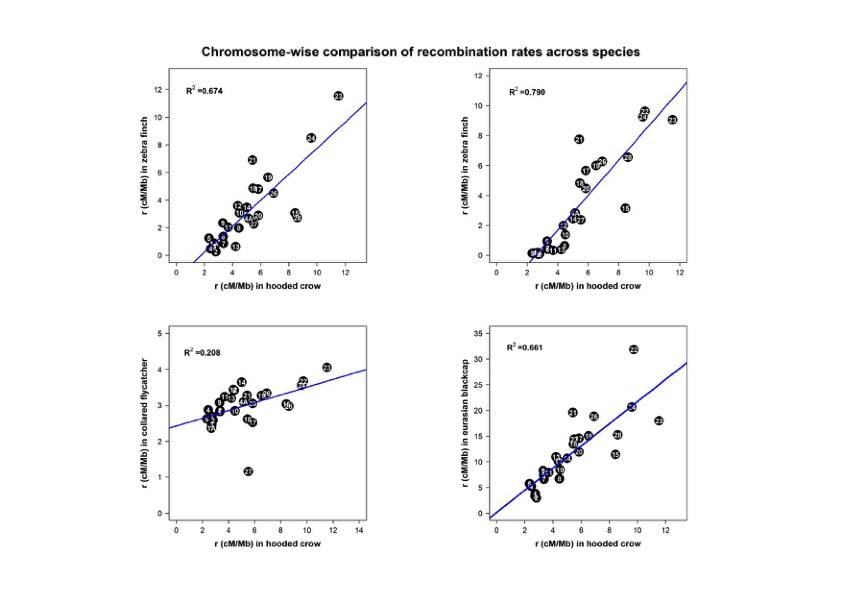
**Supplementary Figure 6 | Correlation of chromosome-level recombination rates of carrion crows with zebra finch, collared flycatcher and blackcap.** For estimates see methods.

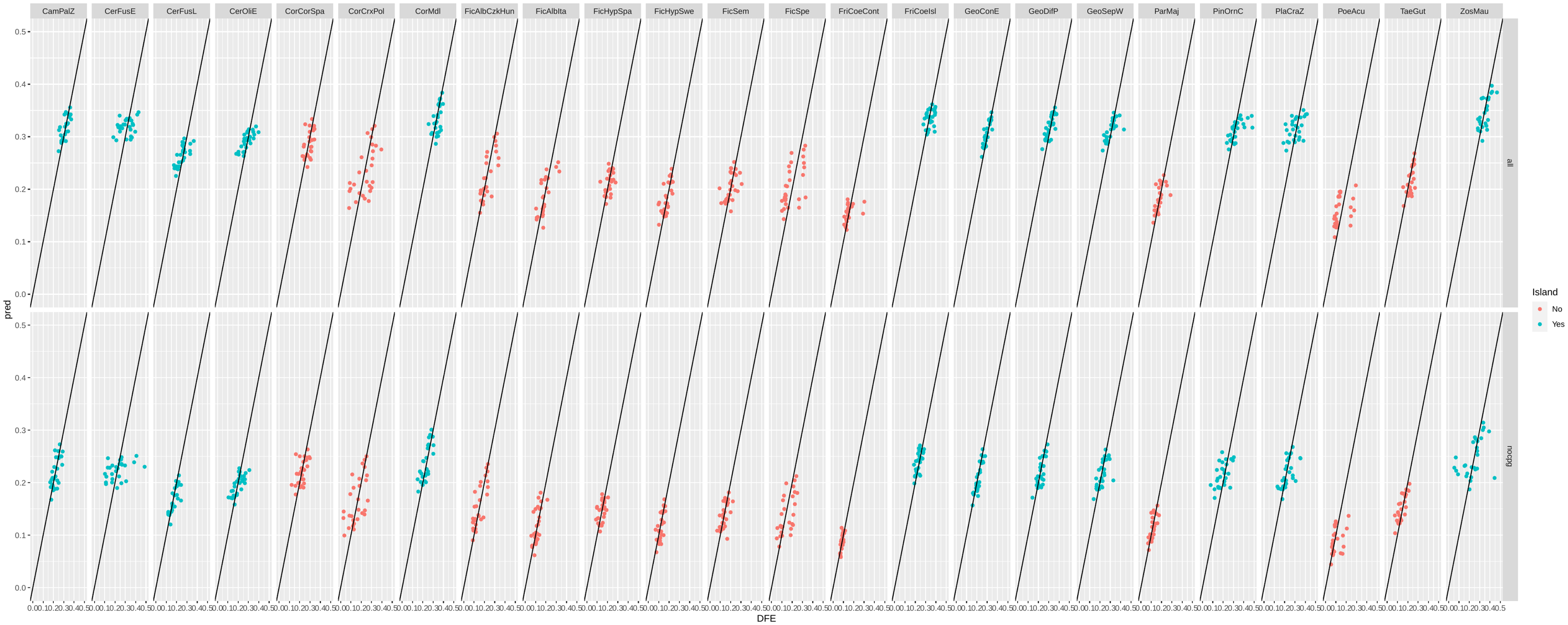

**Supplementary Figure 7 | Model fit.** Relationship between estimates of the proportion of mildly deleterious mutations (*N_e_s* ∈ (-1;0]; DFE) and values predicted (pred) from the best fitting model including recombination rate, effective population size and the effect of biased gene conversion and their interactions (see **Table 1**). Each column represents one of the 24 populations. The upper row shows estimates for all mutations, the lower row excludes CpG-prone sites and weak-to-strong mutations [A,T]->[C,G] subject to biased gene conversion. Colors indicate whether a species is restricted to an island (blue) or has a wider mainland distribution (red). Population abbreviations can be found in **Supplementary Table 2.**

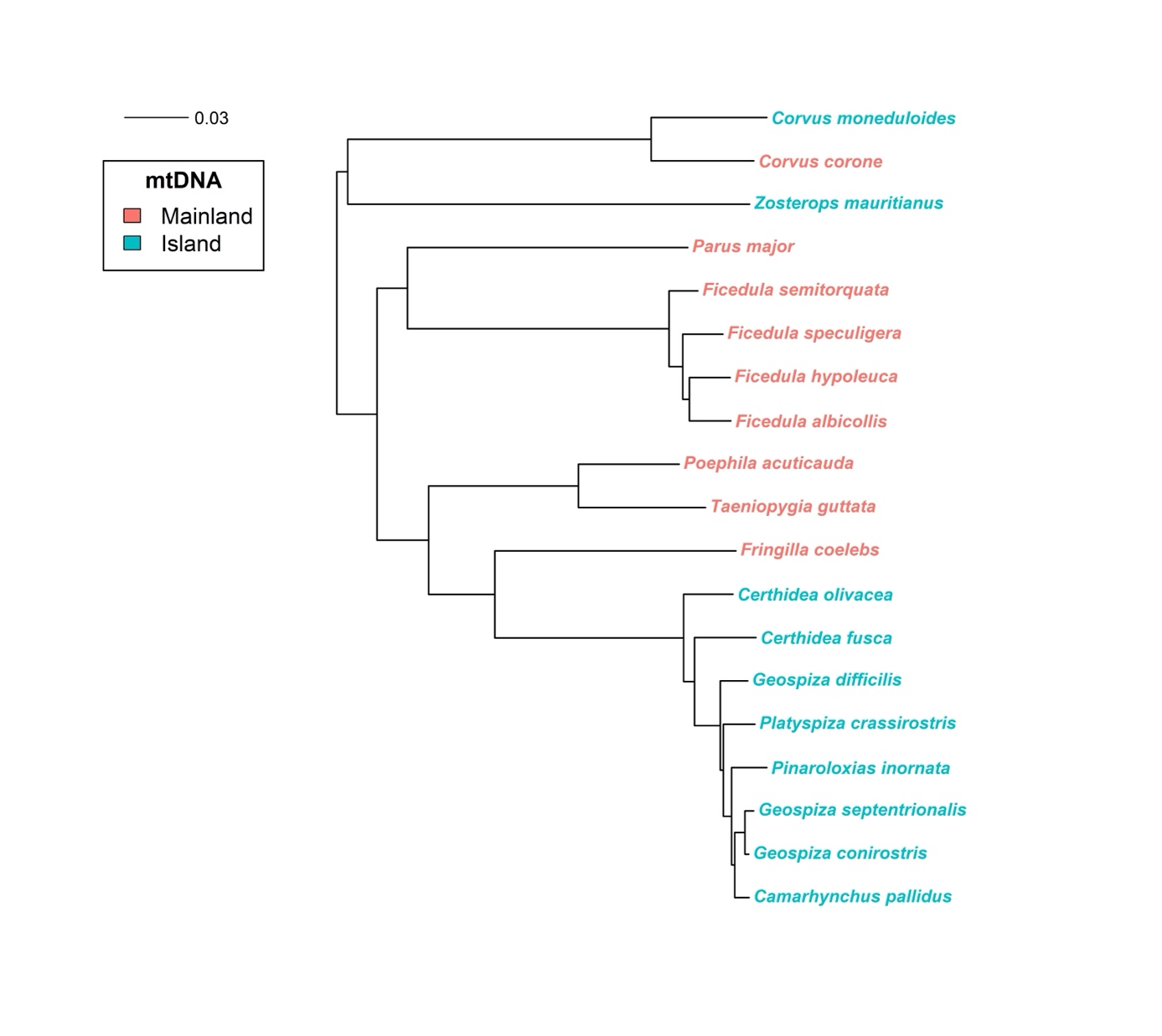

**Supplementary Figure 8 |** **Mitochondrial phylogeny of the 19 species sampled in this study.** The tree was built using protein-coding genes under a GTR+GAMMA+I substitution model and further used as backbone to control for phylogenetic effects.

**Supplementary Tables**

All tables are provided as a separate .xlsx files.

**Supplementary Table 1 | Metainformation on samples and sequencing data.**

***Worksheet 1:*** ***Metadata*** *including species name, subspecies/population information and their abbreviation, literature reference for the data set, the sequencing read archive (SRA) accession number for each sample, original sample ID and alternate sample ID, reference to the genome used for read mapping and an indication whether a sample survived the various filters to be included in the final analysis.*

***Worksheet 2:*** ***Bibliography*** *for the citations of the data set.*

**Supplementary Table 2 | Summary statistics of the data set used for the analyses**. Shown is the number of individuals per population, the number of segregating sites (SNPs) separated by chromosome and selective category (sel: non-synonymous; neutral: 4-fold degenerate), as well as the number of all sites including invariant sites.

***Worksheet 1 (all sites):*** *Summary statistics for all sites.*

***Worksheet 2 (no gBGC):*** *Summary statistics for the subset of sites excluding the effect of biased gene conversion (CpG-prone sites and weak-to-strong mutations).*

**Supplementary Table 3 |** **Summary of statistical models.**

***Worksheet 1*** *List of abbreviations*

***Worksheet 2 (RandomEffects):*** *Statistics for model selection of random effects*

***Worksheet 3 (FixedEffects):*** *Statistics for model selection of fixed effects*

***Worksheet 4*** *Results from the Bayesian model taking phylogenetic dependenc into account*
